# Systems-level analysis identifies IRF6 as an inhibitor of epithelial-mesenchymal transition

**DOI:** 10.64898/2026.01.31.702311

**Authors:** Ayalur Raghu Subbalakshmi, Aditya Agrawal, Shibjyoti Debnath, Kishore Hari, Sarthak Sahoo, Jason A Somarelli, Mohit Kumar Jolly

## Abstract

**Background:** Epithelial-mesenchymal transition (EMT) and its reverse process Mesenchymal-Epithelial Transition (MET) are crucial during metastasis and therapy resistance. While the dynamics and master regulators of EMT are well-studied, the transcription factors that can prevent EMT or promote MET are relatively less understood.

**Results:** Here, by integrating bulk and spatial transcriptomic data analysis from cell lines and patient samples, with mechanism-based dynamical modelling, we identify IRF6 as a factor that strongly associates with an epithelial phenotype and is often inhibited during EMT. *In vitro* experiments in multiple cancer cell lines demonstrate the progression to a mesenchymal phenotype upon IRF6 knock-down, suggesting a role as an inhibitor of EMT. Finally, we observe that IRF6 expression levels correlates with worse patient survival in a subset of solid tumour types.

**Conclusion:** Our integrated computational-experimental systems-level analysis suggests that IRF6 is frequently downregulated during EMT and can also prevent the progression towards a complete EMT, underscoring its role as an MET stabilizing factor.

## Introduction

Cancer remains a major disease burden and a global health challenge. A vast majority of cancer-related deaths are caused by metastasis (1). Metastasis is very inefficient and has extremely high attrition rates (> 99.5%), but a few disseminated cancer cells that eventually lead to colonization achieve that through phenotypic plasticity – their ability to adapt reversibly and quickly to their environments, without changing their DNA sequences (2). Phenotypic plasticity also enables cells to evade multiple therapies, including targeted therapy and immunotherapy (3–5). Thus, elucidating the key drives of phenotypic plasticity is essential to eventually develop therapeutic strategies to mitigate metastasis and drug resistance.

A very well-studied example of phenotypic plasticity is the set of reversible processes of epithelial to mesenchymal transition (EMT) and its reverse – mesenchymal to epithelial transition (MET). During a partial or full EMT or MET, cells can transition back and forth between epithelial (E), mesenchymal (M) and hybrid E/M phenotypes (6). The signaling pathways (TGF-β, EGF, TNF) (7,8) and downstream transcription factors (TFs) driving EMT – ZEB1, ZEB2, SNAI1, SNAI2, TWIST (9–12) – have been extensively characterized. Recent studies have also suggested signaling pathways and TFs that can stabilize one or more hybrid E/M phenotypes such as NFATc and NRF2 (13–16). Similarly, MET-inducing microRNAs such as miR-200 family and MET-TFs such as OVOL1 and GRHL2 have been reported (17,18) that form a mutually inhibitory loop with EMT-TFs such as ZEB1 (19–22). Although extensive time-course bulk and single-cell transcriptomic data on EMT progression has been obtained (7,23), the dynamics of MET is less well understood, and has been mostly studied only as withdrawal of EMT-inducing signal rather a direct induction of MET itself (8). Cells undergoing MET due to withdrawal of the EMT-inducing signal, after having undergone EMT, may not return to the same epithelial state prior to EMT induction – a phenomenon known as hysteresis – as observed at transcriptomic, proteomic and epigenetic cells, suggesting a cellular memory of EMT (24–28). Also, cells ‘locked’ in ‘extremely mesenchymal’ states have limited metastatic ability (29,30) relative to plastic hybrid E/M states. Thus, mapping the dynamics and drivers of MET is crucial to curb tumor aggressiveness.

Interferon regulatory factor 6 (IRF6) belongs to interferon (IFN) regulatory transcription factor (IRF) family that has 9 members (IRF1-IRF9). Unlike other IRF family members that are involved in immune responses, IRF6 does not modulate IFN gene expression. Instead, it is crucial for craniofacial morphogenesis (31) and in development of other tissues derived from the ectoderm (32). IRF6-deficient mice have compromised terminally differentiated epidermal layers, leading to abnormal skin, limb and craniofacial development (33). Recent studies have reported the role of IRF6 in EMT in cancer; among breast cancer cell lines, reduced IRF6 mRNA and protein levels were observed in more mesenchymal MDA-MB-231 and MCF10A cells as compared to epithelial MCF7 cells (34). Similarly, IRF6 downregulation correlates with tumor invasive and differentiation status in squamous cell carcinoma (35). On the other hand, over-expression of IRF6 led to upregulated levels of epithelial players (E-cadherin, desmoplakin) and reduced levels of mesenchymal markers (Vimentin, N-cadherin) in SW480 colorectal cancer cells (36), and in S18 and 5-8F nasopharyngeal cancer cells (37). Reciprocally, EMT induction in pancreatic cancer silenced IRF6 at epigenetic and transcriptional levels (38). Together, these studies suggest IRF6 as a putative MET-TF.

Here, we analyze bulk and spatial transcriptomic datasets from cancer cell lines and patient samples to identify novel factors associated with MET. These analyses pinpointed IRF6 to be strongly associated with an epithelial phenotype and its downregulation upon induction of EMT. We next develop a mechanism-based mathematical model by integrating the reported interactions among IRF6 and key EMT/MET regulators. Simulations suggest that knockdown of IRF6 can induce EMT, which were then experimentally validated in SW480, MCF7 and A431 cells. Patient survival data analysis pinpointed a tissue-specific association between IRF6 levels and survival. Together, our work proposes IRF6 as an inhibitor of EMT and a prospective MET-TF.

## Results

### Association of IRF6 with an epithelial phenotype

We first examined the association between IRF6 expression levels and the pan-cancer epithelial and mesenchymal gene expression programs (39). We calculated single-sample gene expression enrichment analysis (ssGSEA) scores (40) for the epithelial and mesenchymal gene lists and correlated them with expression levels of all genes, using the cohort of Cancer Cell Line Encyclopedia (CCLE) (41) that contains over than 900 cell lines across cancer types. As expected, canonical epithelial genes such as CDH1, OVOL2, GRHL2 and ELF3 correlated positively with ssGSEA-based epithelial score and negatively with ssGSEA-based mesenchymal scores. Conversely, many mesenchymal genes such as ZEB1, SNAI1, SNAI2 and VIM correlated positively with ssGSEA-based mesenchymal score and negatively with the epithelial scores. IRF6 was found to be clustered along with the epithelial genes in the fourth quadrant, unlike other IRFs (**Fig 1A**). In the CCLE cohort, IRF6 was the only IRF family member that showed a strong positive correlation with MET-T TFs such as ELF3, KLF4, OVOL1, OVOL2 and GRHL2 and a negative correlation with EMT-TFs such as ZEB1 and ZEB2 (**Fig 1B**). To further support these observations, we analyzed data from TCGA across cancer types and observed that IRF6 expression levels were negatively correlated with the ssGSEA-mesenchymal score and positively correlated with the ssGSEA-epithelial scores in multiple cancer types such as lung, ovarian, colon and breast cancers (**Fig 1C**), further indicating that IRF6 associates with an epithelial phenotype.

**Figure 1.**
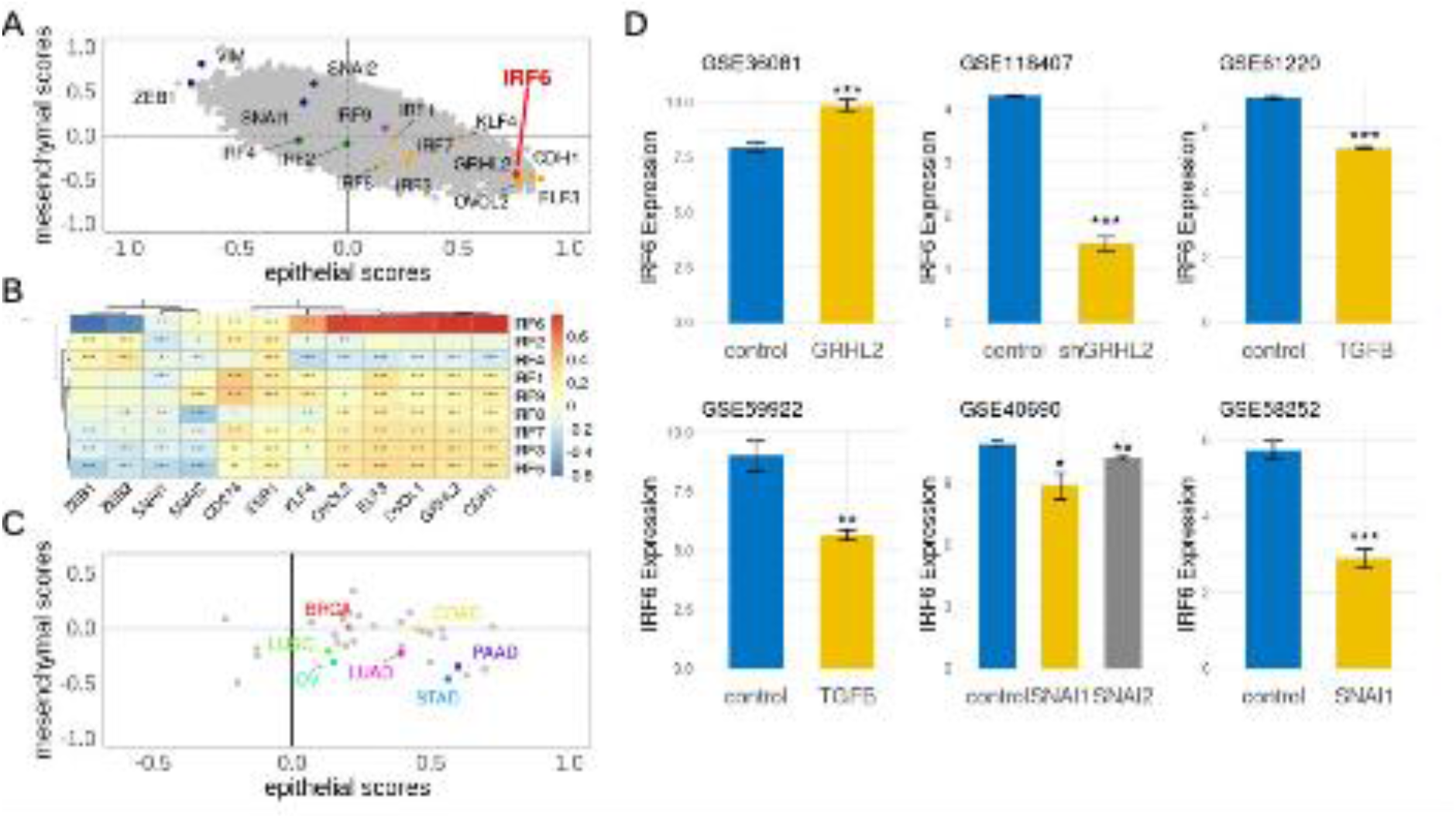
IRF6 expression levels correlate positively with an epithelial phenotype. A) Scatterplot showing the correlation coefficient values of individual gene expression levels with epithelial and mesenchymal scores in CCLE cohorts. Mesenchymal genes VIM, ZEB1, SNAI1 and SNAI2 are represented in blue and epithelial genes GRHL2, OVOL2, KLF4, CDH1 and ELF3 are represented in orange. IRF2 and IRF4 are represented in green and IRF9 in purple. IRF6 is represented in red. B) Correlation of different IRF1-9 expression levels with other epithelial and mesenchymal genes in CCLE cohort. C) Correlation of IRF6 expression levels with epithelial and mesenchymal scores in different TCGA cancer types. D) Variation in IRF6 expression levels in multiple GEO datasets upon EMT and/or MET induction: i) GSE36081 ii) GSE118407 iii) GSE61220 iv) GSE59922 v) GSE40690 vi) GSE58252

Next, we investigated whether the levels of IRF6 varied as a function of EMT/MET induction across multiple, independent data sets. We observed that induction of MET through GRHL2 overexpression in HMLE cells that express a Twist-ER fusion (a model of inducible EMT) led to increased IRF6 levels (GSE36081) (42) (**Fig 1D, i**). Conversely, induction of EMT in OVCA4209 cells (GSE118407) (43) by silencing GRHL2 led to reduced expression of IRF6 (**Fig 1D, ii**). Similarly, induction of EMT by TGFβ suppressed IRF6 levels in airway epithelial cells (GSE61220) (44) and in mouse mammary EpRas cells (GSE59922) (45) (**Fig 1D, iii-iv**). Further, induction of EMT through overexpression of SNAI1 or SNAI2 in human mammary epithelial cells (GSE40690) (46), or via SNAI1 in MCF7 cells (GSE58252) (47) consistently showed reduced IRF6 expression (**Fig 1D, v-vi**). These above-mentioned patterns are consistent with IRF6 downregulation in the mesenchymal subpopulations of prostate cancer epithelial cells PC3 and its upregulation upon lentiviral ZEB1-shRNA vector treatment (48). Moreover, these patterns are specific to IRF6 and were not observed as coherently in other IRF6 family members (IRF1-IRF5, IRF7-IRF9) (**Fig S1A-H**). Together, these observations suggest that downregulation of IRF6 is a salient feature of EMT induction.

### IRF6-associated signature marks epithelial-like states in breast cancer and is inversely associated with mesenchymal programs

To further delineate the regulatory features underlying EMT states in breast cancer, we investigated the role of IRF6 using single-cell RNA-sequencing data derived from ER+ and triple-negative breast cancer (TNBC) patients (49). We sought to understand how the IRF6-associated transcriptional program (50) corresponds with established breast cancer-specific epithelial, mesenchymal, and partial EMT (pEMT) gene signatures (51). Cells from ER+ tumors (shown in black) and TNBC tumors (shown in red) display distinct expression patterns for the IRF6-associated signature (**Fig 2A-C**).

**Figure 2.**
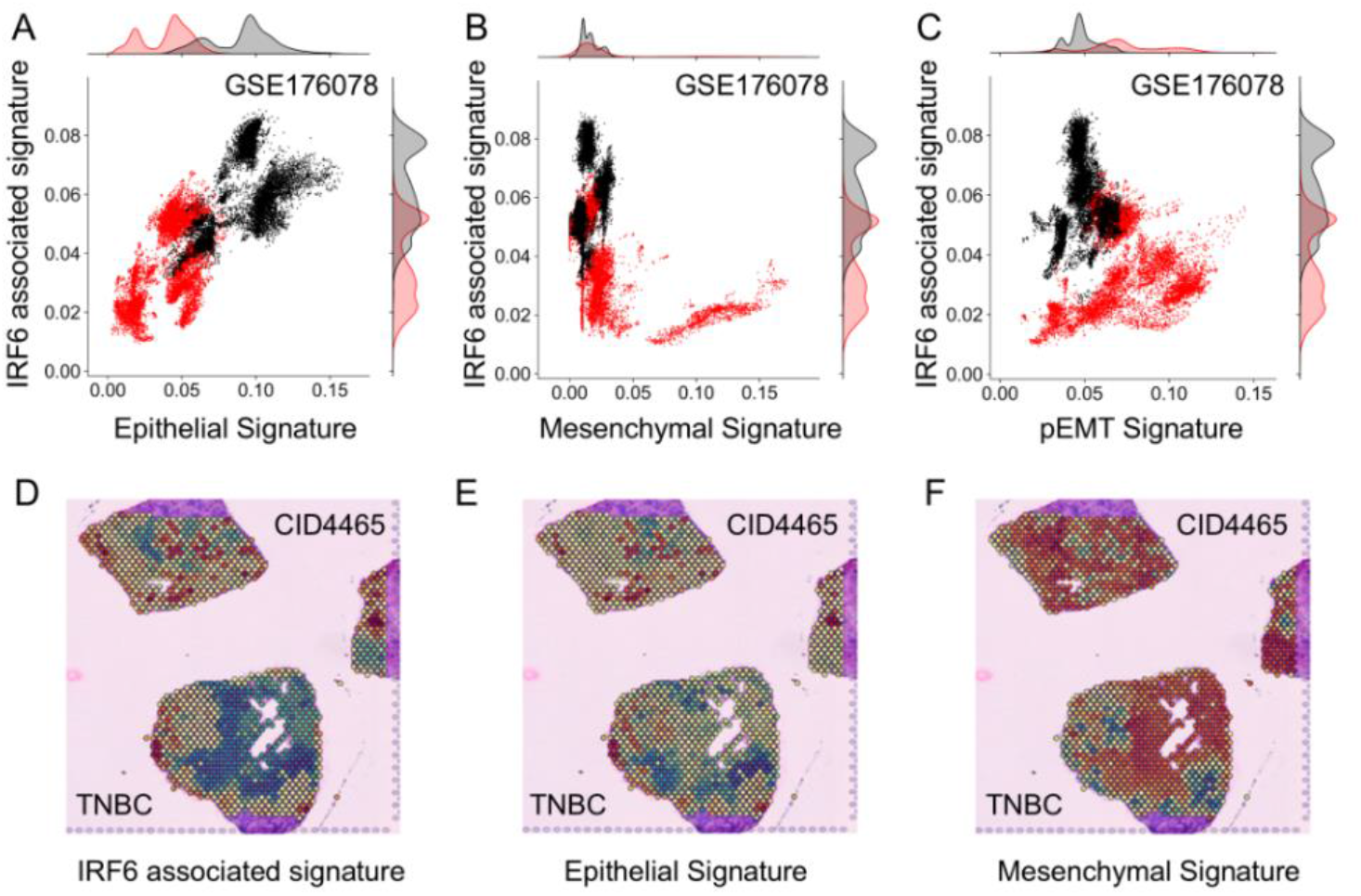
IRF6-associated signature is co-expressed with epithelial signatures and is anti-correlated with mesenchymal signatures across breast cancer cells. A) Scatterplot showing the co-expression of IRF6-associated and breast cancer-specific epithelial signatures in single-cell RNA-seq data from ER+ (black) and TNBC (red) breast cancer patients. Spearman correlation: r = 0.72, p-val <0.01. B) Scatterplot showing the inverse relationship between IRF6-associated and breast cancer-specific mesenchymal signature scores in the same dataset. Spearman correlation: r = -0.11, p-val <0.01. C) Scatterplot of IRF6-associated versus breast cancer-specific partial EMT (pEMT) signature. Spearman correlation: r = -0.29, p-val <0.01. Spatial plots of D) IRF6 associated signature, E) breast cancer-specific epithelial signature and (F) breast cancer specific mesenchymal signature activity in TNBC tumor section (CID4465).

Strikingly, we observed that ER+ breast cancer cells exhibited consistently high activity score for IRF6-associated signature, which co-occurred with high expression of a breast cancer-specific epithelial program (**Fig 2A**). These same cells concurrently showed lower expression of the breast cancer-specific mesenchymal signature (**Fig 2B**). While the association between IRF6 and a breast cancer-specific pEMT signature was mildly positive (**Fig 2C**), it was not uniformly distributed across the cell populations and may reflect intermediate epithelial-hybrid states rather than the direct stabilization of pEMT by IRF6. Similarly, breast cancer cells from TNBC tumors that expressed higher levels of epithelial signature also expressed higher IRF6 associated signature (**Fig 2A**) while TNBC cells that expressed the highest levels of mesenchymal signature expressed the least IRF6 associated signatures (**Fig 2B**).

To determine whether similar trends were observed at a tissue level, we analyzed spatial transcriptomics data from TNBC breast cancer patients (CID4465, CID44971) (52) who are known to express considerable heterogeneity in EMT states (53), to assess the spatial co-localization of the IRF6 signature with epithelial and mesenchymal programs. In tumor sections, regions with high IRF6 signature activity consistently co-localized with areas enriched for the epithelial signature (**Fig 2D-E, S2A-B**) and were mutually exclusive with regions displaying high mesenchymal scores (**Fig 2F, S2C**).

Together, these results provide transcriptomic evidence – both at the single-cell and spatial levels – that the IRF6-associated program preferentially marks epithelial-enriched subpopulations across the breast cancer subtypes and is downregulated in mesenchymal-like states. These results support the hypothesis that IRF6 may function as a MET-promoting transcription factor.

### IRF6 inhibits EMT progression

Next, we assessed the role of IRF6 in mediating EMT dynamics. To do this, we constructed a gene regulatory network (GRN) based on experimental evidence including the interactions of IRF6 with a previously studied core EMT regulatory network (54) (denoted by black dotted rectangle in **Fig 3A**). This network comprised both epithelial factors, such as miR-200, KLF4 and ELF3 and mesenchymal factors, such as ZEB, SNAI1 and SNAI2. IRF6 has been shown to activate epithelial factors such as KLF4 (55), GRHL3 (56) and GRHL2 (57), and is critical for proper localization of E-cadherin to the cell membrane, enabling cell adhesion (58). In turn, IRF6 is activated by two MET-TFs, namely ELF3 (59) and GRHL2(57). Conversely, ZEB1, an EMT-TF, can inhibit IRF6 directly (59). ZEB1 can repress E-cadherin transcriptionally ; reciprocally, E-cadherin can sequester β-catenin to the cell membrane, preventing its nuclear localization and consequent activation of ZEB1, thus effectively inhibiting ZEB1(60,61).

**Figure 3.**
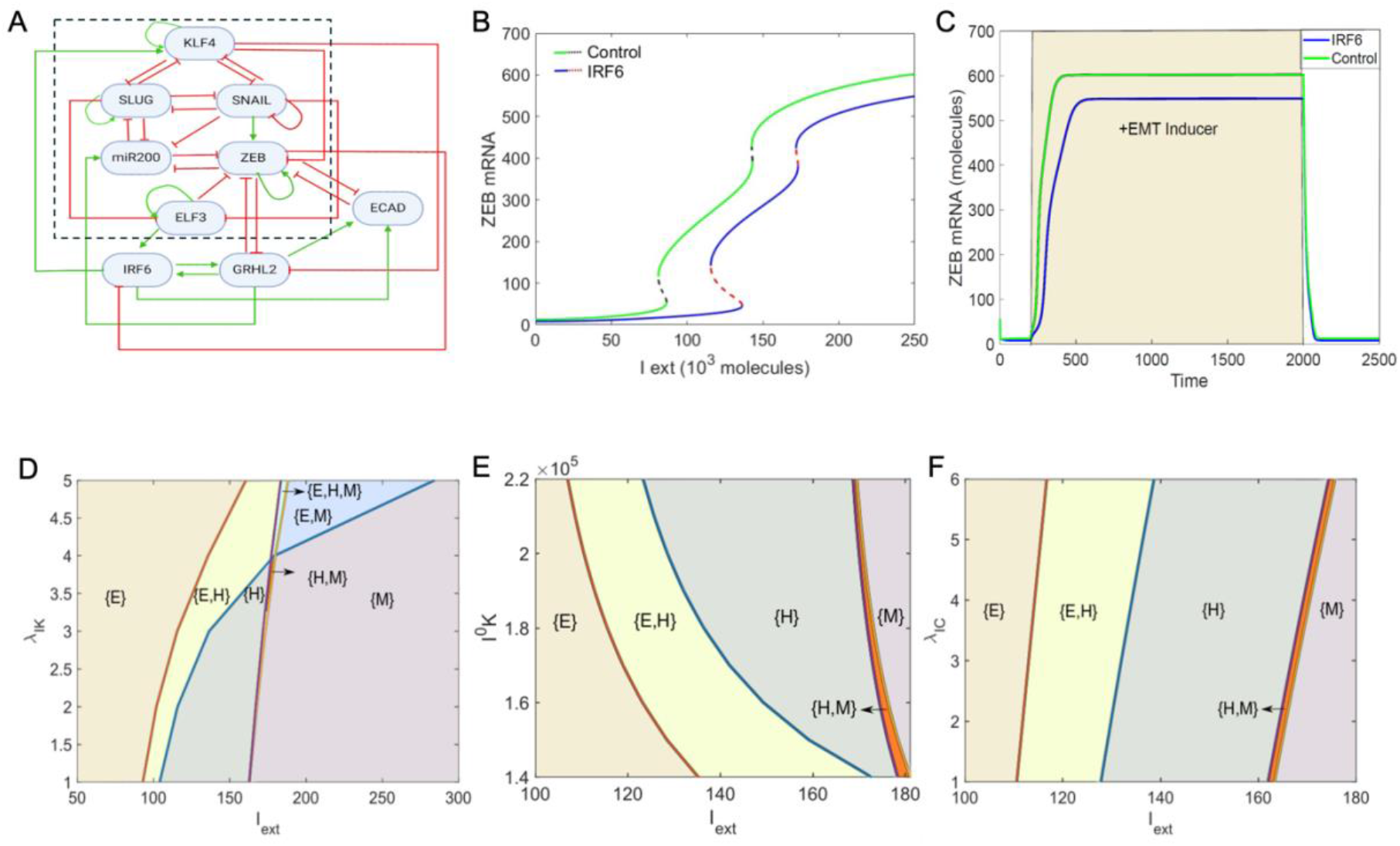
IRF6 inhibits EMT induction. A) Schematic representation of IRF6 coupled with an EMT regulatory network (dotted rectangle) consisting of miR-200, ZEB, SNAIL, SLUG, KLF4 and ELF3. Green arrows denote activation, and red bars indicate inhibition. Solid arrows represent transcriptional regulation; dotted lines represent microRNA-mediated regulation. B) Bifurcation diagram indicating the changing levels of ZEB mRNA levels in response to levels of an external signal (I_ext) level for the coupled EMT–IRF6 circuit (solid blue and dotted red curve) and the core EMT circuit (solid green and dotted black curve). C) Temporal dynamics of ZEB mRNA levels as a response to a high level of an external EMT signal (I_ext = 200,000 molecules) in an epithelial cell-state (yellow-shaded region) for the circuit shown in panel A. D) Phase diagrams for coupled EMT–IRF6 network driven by an external signal (I_ext) for varying strength of activation by IRF6 on KLF4. E) Phase diagrams for coupled EMT–IRF6 network driven by an external signal (I_ext) for varying threshold levels of IRF6 needed for KLF4 activation F) Phase diagramsfor coupled EMT–IRF6 network driven by an external signal (I_ext) for varying strength of activation by IRF6 on ECAD.

Using these established interactions, we developed a mathematical model to determine the effect of IRF6 on EMT progression. First, we plotted a bifurcation diagram of ZEB mRNA levels upon EMT induction by an external signal I_ext_ (**Fig 3B**). We observed that with an increase in I_ext_, the cells transitioned from an epithelial phenotype, as indicated by low levels of ZEB mRNA levels (< 100 molecules) to a hybrid E/M phenotype, indicated by intermediate levels of ZEB mRNA levels (100 < ZEB mRNA <400) and ultimately to a mesenchymal phenotype indicated by high levels of ZEB mRNA molecules (> 450). In the absence of IRF6 (solid green and dotted black curve), we observe that the transitions happen at lower levels of I_ext_, as compared to the circuit where the above-mentioned interactions of IRF6 are included (solid blue and dotted red curve), pointing out that a stronger EMT-inducing signal is needed for the onset of EMT in the presence of IRF6. When we analyzed the temporal dynamics for a fixed value of I_ext_ (**Fig 3C**), we observed that IRF6 delayed EMT and resulted in lower levels of steady state ZEB levels (blue curve) when compared to the control network (green curve). We also calculated the bifurcation diagrams and temporal dynamics for different variants of the network by simulating IRF6, GRHL2 and/or E-cadherin knockdown, which also pointed to a strong role of IRF6 in impacting EMT (**Fig S3A, i-ii**).

We next estimated the extent to which the IRF6 mediated impact on EMT was dependent on its interactions with the EMT circuit. When the strength of activation of KLF4 by IRF6 was increased, we observed an expansion of the {E} region (epithelial phenotype) and a reduced {M} region (mesenchymal phenotype (**Fig 3D**). Similarly, when we varied the threshold levels of IRF6 needed to activate KLF4 (**Fig 3E**), we observed that at lower threshold levels (i.e. effective stronger activation), the {E} phase was favored; however, as the threshold level increased, the {M} phase was favored. Consistent observations were made when we varied the strength of activation on CDH1 by IRF6 (**Fig 3F**): the stronger the activation, the more likely an epithelial phenotype. To further evaluate how specific our model predictions are to the chosen values of kinetic parameters, we performed sensitivity analysis by varying the numerical values of these parameters by ± 10% one by one and measured the changes in the range of I_ext_ values enabling a hybrid E/M state in the bifurcation diagram. Except for a few parameters, the percentage change was found to be less than 15% (**Fig S3B**). This analysis underscores the notion that the ability of IRF6 to inhibit EMT is robust to small variations in parameter values.

### Knockdown of IRF6 can drive EMT

To further analyze the role of IRF6 in shaping the phenotypic landscape, we employed a computational framework known as RACIPE (62) to simulate the emergent network dynamics of the GRN. RACIPE takes the structure of a GRN as input and converts it into a system of coupled ordinary differential equations (ODEs). It then solves these ODEs across a broad spectrum of biologically relevant parameter values and produces as output the steady-state gene expression patterns (phenotypes) that the GRN can produce. We plotted the steady-state patterns as a heatmap, where hierarchical clustering revealed two clusters: one cluster consisted of high ZEB and SLUG levels and low levels of epithelial markers (miR200, GRHL2, CDH1, KLF4, ELF3 and IRF6) and the other cluster comprises of low levels of ZEB and SLUG but high levels of epithelial markers (red and green vertical bars in **Fig 4A**). Next, we performed *in silico* overexpression of IRF6 that led to a decreased frequency of the mesenchymal phenotype accompanied by an increase in that of the epithelial phenotype. A comparison with *in silico* overexpression of ELF3 and KLF4 exemplified a stronger role of IRF6 in promoting MET (**Fig. 4B, i**). Similarly, *in silico* knockdown of IRF6 resulted in a decreased frequency of the epithelial phenotype (**Fig 4B, ii**), indicating that IRF-KD can induce EMT.

**Figure 4.**
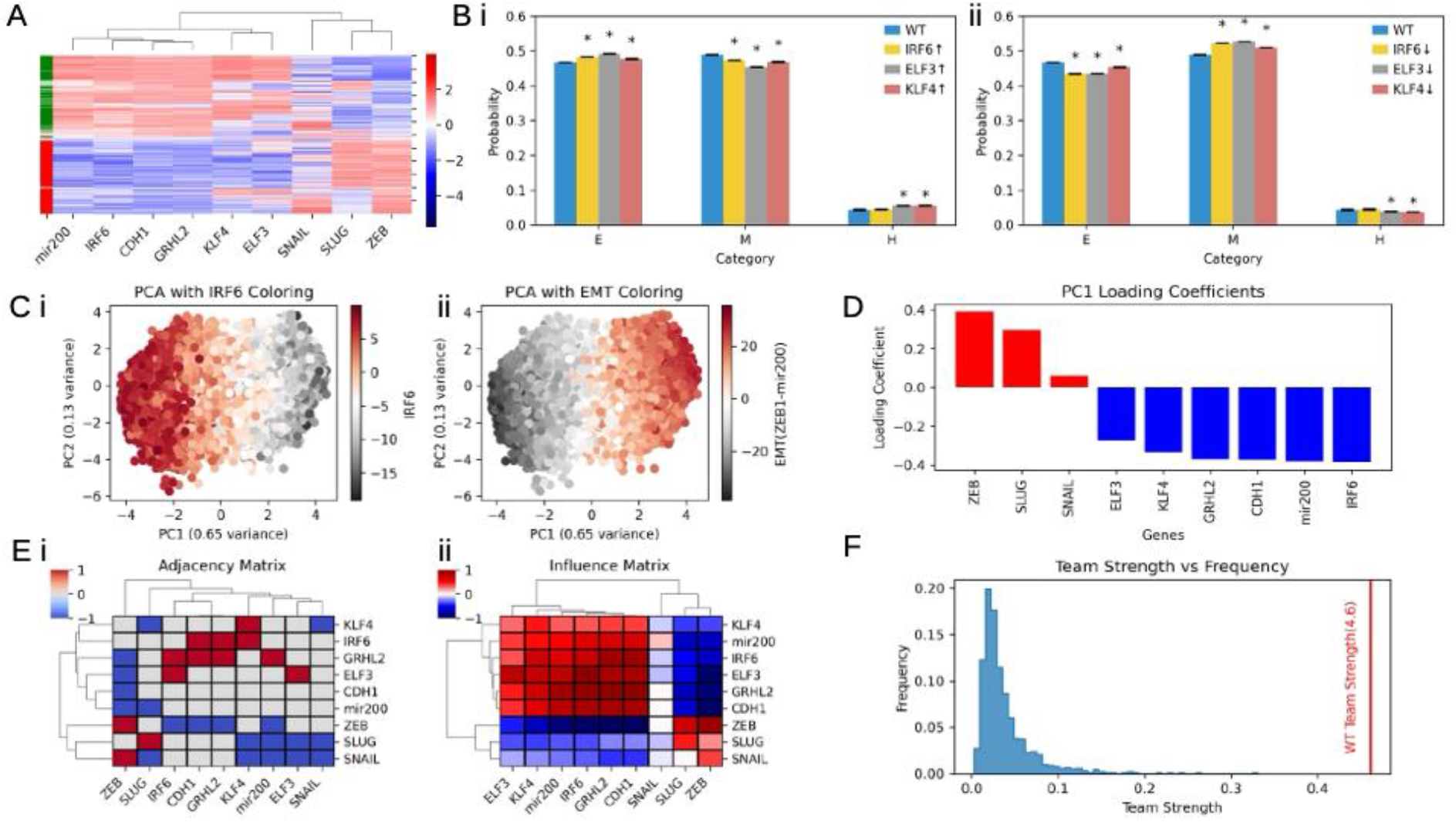
IRF6 belongs to the team of factors promoting an epithelial phenotype. A) Heatmaps depicting the steady state values of RACIPE simulations of the GRN shown in Fig 3A. B) Steady-state phenotype frequency upon i) over-expression of IRF6, ELF3 and KLF4 ii) down-expression of IRF6, ELF3 and KLF4, as per RACIPE. C) PCA scatter plot of all steady states of RACIPE colored by, i) IRF6, and ii) EMT score (= ZEB – miR200), where principal component 1 (PC1) explains 65% of the variance. D) Bar plot showing the PC1 loading coefficients of different genes in the network. E) i) Adjacency matrix for the network shown in Fig 3A. ii) Influence matrix for path length = 10 for network shown in Fig 3A, showing two teams – an EMT-promoting (ZEB, SNAIL, SLUG) and an EMT-inhibiting (KLF4, miR-200, IRF6, ELF3, GRHL2, CDH1) one. F) Histogram of the team strength of 1000 random networks generated. The red line indicates the team strength of the wild-type (WT) network.

Next, we performed a principal component analysis (PCA) of the z-normalized steady-state solutions. We observed that 65% variance was captured by the PC1 axis. The PCA plot revealed two main clusters along the PC1 axis: one with reduced IRF6 levels and higher EMT scores (= ZEB1 – miR-200) and the other one with higher IRF6 levels and reduced EMT scores (**Fig 4C, i-ii**). These trends indicate that across the parameter sets considered, this network can recapitulate phenotypic heterogeneity along the epithelial-mesenchymal (E-M) spectrum. To quantify the contributions of each gene to PC1, we examined the PC1 loading coefficients (**Fig 4D**). Nodes with positive coefficients corresponded to mesenchymal markers (ZEB1, SLUG and SNAIL), while those with negative coefficients corresponded to epithelial markers (ELF3, KLF4, GRHL2, CDH1, miR-200 and IRF6), further supporting the notion that PC1 variance captured E-M heterogeneity. IRF6 had the highest absolute value of loading coefficients among all the nodes, highlighting its central role in governing phenotypic variation along the E-M axis.

Further, we represented the GRN in the form of an adjacency matrix that represented direct interactions in the GRN. Each element in this matrix can take three values (+1 for an activation link, -1 for an inhibition link, and 0 for no direct link). Each row represented the source node, and each column denoted a target node (**Fig 4E, i**). Given that nodes in a GRN can influence one another through indirect paths of varying lengths as well, we derived an influence matrix by summing the adjacency matrix raised to powers from 1 to 8, thus capturing the cumulative effect of paths up to length 8 (**Fig 4E, ii**). Each element in this influence matrix captures the net effect or influence a source node (row) has over a target node (column) over multiple paths of varying lengths, with higher weightage given to shorter path lengths (63). The influence matrix revealed two mutually inhibitory “teams” of nodes such that members within a team effectively activated each other, while those across the two teams effectively inhibited one another. One team was composed of ZEB, SNAIL and SLUG, while the other consisting of KLF4, miR-200, ELF3, GRHL2, CDH1 and IRF6, mirroring the trend observed in PC1 loading coefficients. Thus, network topology analysis also supported that IRF6 belonged to the team of MET drivers or EMT inhibitors, in mutual antagonism to the team of molecular players driving EMT.

To quantify the coordination among nodes within and across these teams, we calculated a metric called ‘team strength’ (defined on a scale of 0 to 1) to be 0.45. The higher the ‘team strength’, the more robust the antagonism across teams and mutual reinforcement within teams (63). To assess how specific this teams structure is to this GRN, we generated 1000 random networks of the same size, by shuffling the regulatory links in the GRN, while preserving degree distributions. We calculated the ‘team strength’ and PC1 variance for all 1,000 networks and found that the wild-type (WT) GRN had the highest ‘team strength’ and PC1 variance (**Fig 4F, S4A-B**), suggesting that the two-team structure is a non-random, functionally selected feature of this GRN. Also, the random networks exhibited no trend in their loading coefficients (**Fig S4C**) as was identified for the WT network, further reinforcing the observation that the WT network contains a strong functional antagonism between EMT and MET drivers, of which IRF6 is a part of the epithelial team.

We observed similar results when we performed Boolean simulations of this network. Boolean framework allows for only two states of a node in a network: ON (high or +1) or OFF (low or -1). We adopted an Ising model formalism here, where each edge in the network has equal weightage, and the state a node takes in each time-step depends on a simple majority rule among the incoming edges: if the number of inhibitory edges dominate, the node is turned OFF, and if excitatory edges dominate, it is turned ON (64). For 10,000 random initial conditions that we simulated across, 74% of cell-state trajectories converged to two steady states (phenotypes): epithelial (E) and mesenchymal (M) defined here based on the difference in mean expression levels of the mesenchymal (SNAIL, ZEB and SLUG) and epithelial (miR-200, ELF3, KLF4, ECAD and GRHL2) that showed a strongly mutually exclusive pattern. In most steady-state solutions where ZEB1 was high, we observed high levels of SNAIL and SLUG as well, and conversely in most cases where miR-200 levels were high, various epithelial markers were high (ELF3, IRF6, KLF4, GRHL2, ECAD) (**Fig 5A**). We then performed *in silico* overexpression (OE) and knockdown (KD) experiments for MET factors (GRHL2, ELF3, KLF4 and IRF6) one at a time, with SNAIL OE or KD as a negative control. As expected, we found that over-expressing MET factors increased the frequency of epithelial phenotype and reduced that of mesenchymal one, whereas their simulated knockdown had the opposite effect (**Fig 5B, i-ii**).

**Figure 5.**
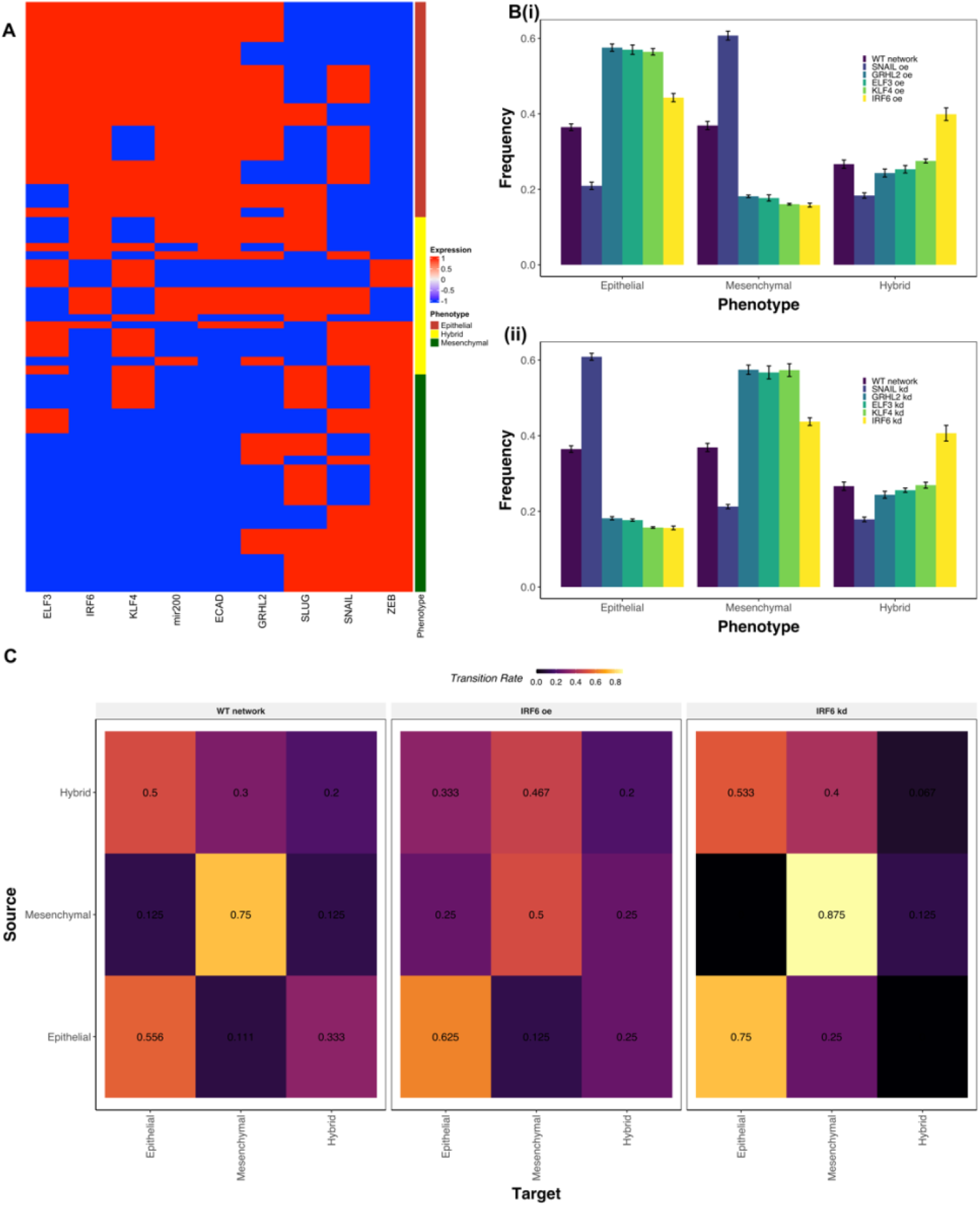
Analysis of the role of IRF6 in EMT/MET using Boolean simulations. A) Heatmap depicting the steady states obtained from simulating the network in Fig 3A using Boolean formalism starting from 100000 initial conditions. Each row corresponds to a steady state and each column corresponds to the expression level of each node across all steady states. High expression (+1) is shown in red and low expression (-1) in blue. A strip adjacent to the heatmap represents the phenotype of each steady state (defined based on ZEB and miR-300). B) Bar plots depicting the change in the frequency of epithelial, mesenchymal, and hybrid phenotype for WT and overexpression (oe - i) or down expression (kd - ii) of SNAIL (negative control), GRHL2, ELF3, KLF4 and IRF6. C) Heatmap depicting the transition rates of states from the source phenotype to that of the target phenotype for WT network, IRF6 overexpression, and down expression cases. The color bar shows the transition rates defined on a scale of 0 to 1. Rows shows the source (initial) phenotype prior to perturbation, columns show the target phenotype attained. Numbers (the three transition rates from an initial state) in each row total to 1.

We attempted to understand the effect of IRF6 and other MET factors on the rate of transition between E, M, and hybrid E/M phenotypes. Thus, we exposed the steady states of the network to transient perturbations, comprised of inverting the signs from -1 to +1 or *vice versa*) for one-third of the randomly selected nodes. We then allowed the dynamics to reach a steady state and calculated the frequency with which different phenotypes were attained. For the WT network, we observed a higher likelihood of E and M phenotypes to be retained, as compared to hybrid E/M, based on transition rates starting from E, M and hybrid E/M as the initial conditions (**Fig 5C, right**). This behavior is reminiscent of EMT GRNs reported earlier (63,64). Upon simulated over-expression of IRF6, we observed twice the likelihood of acquiring an E state when the M state is perturbed transiently (0.25 vs. 0.125; compared **Fig 5C, left** with **Fig 5C, middle**). Consequently, the likelihood of retaining an M state decreases (0.75 vs. 0.5) and that of retaining an E state increases modestly (0.62 vs. 0.56). Upon *in silico* knockdown of IRF6, we noticed a higher likelihood of retaining an M state when perturbed (0.875 vs. 0.75) and more than twice the likelihood of an E state to switch to an M state when perturbed (0.25 vs. 0.11; compare **Fig 5C, left** with **Fig 5C, right**). Together, these observations indicate that knockdown of IRF6 increases the likelihood of cells to attain or maintain a mesenchymal state, while its overexpression can contribute to gatekeeping an epithelial phenotype and/or induce MET.

To test these *in silico* predictions from ODE models and the Boolean framework, we performed siRNA-mediated knockdown IRF6 in three cancer cell lines: SW480 (human colon adenocarcinoma), A431 (human epidermoid carcinoma) and MCF-7 (human breast cancer). We observed significant knockdown of IRF6 at the mRNA level by >90%, and a concomitant downregulation of E-cadherin and OVOL1 mRNAs, and upregulation of the EMT factor, ZEB1, in SW480 cells (**Fig 6A, i**). Consistent with this, IRF6 knockdown also led to downregulation of E-cadherin protein and an increase in ZEB1 protein levels in SW480 cells. (**Fig 6A, ii**). A431 and MCF-7 cells exhibited similar patterns upon IRF6 knockdown, with loss of E-cadherin and OVOL1 mRNAs and upregulation of ZEB1 mRNA (**Fig 6B-C**). These experimental observations validate our prediction of IRF6 being an EMT inhibitor.

**Figure 6.**
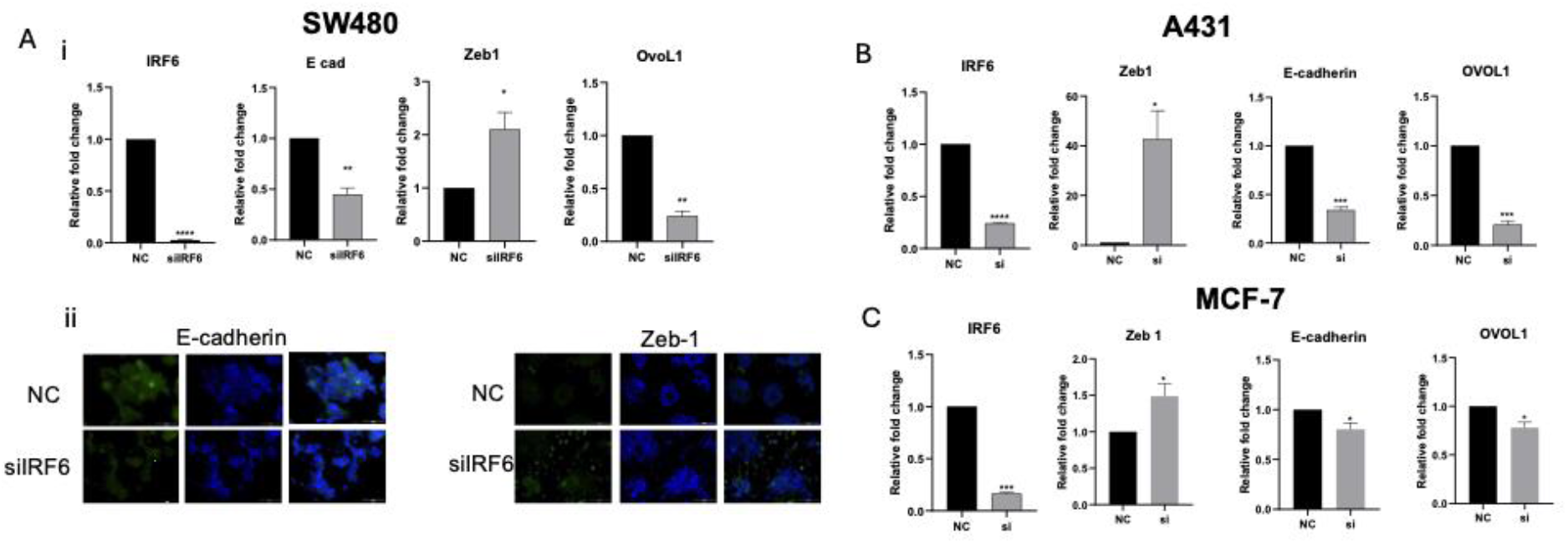
IRF6 knockdown induces gene expression changes consistent with EMT. A) SW480 cells were transfected with IRF6 siRNA, and the relative expression of the EMT markers E-cadherin, ZEB1, and OVOL1 was assessed compared with a non-silencing siRNA control (NC) using RT-qPCR (i) and immunofluorescence microscopy (ii). B-C) A431 and MCF-7 cells were similarly analyzed for the indicated EMT markers following IRF6 siRNA transfection. Results are shown as mean ± SEM. Student’s t-test was used for analysis (two-tailed; ^*^p < 0.05, ^**^p < 0.01, ^***^p < 0.001 vs non-treated, ns not significant).

### IRF6 levels correlated with patient survival in a tissue-specific manner

The role of IRF6 in restricting EMT motivated us to examine its clinical relevance as a biomarker. In gastric cancer, higher IRF6 expression levels associated with better overall survival (GSE62254, GSE22377), and progression-free survival (**Fig 7A-B, S5A**) (65,66). In two ovarian cancer datasets too, we found that higher IRF6 levels were associated with better overall survival, progression-free survival (GSE63885, GSE14764) (**Fig 7C-D, S5B)** (67) and post-progression-free survival **(Fig S5C-D**) (68). However, the trend was opposite in colon cancer, where we noted that higher IRF6 levels correlated with worse overall patient survival and relapse-free survival in many independent datasets: GSE41258 (69), GSE17538 (70) and GSE39582 (71) (**Fig 7E-F, S5E-F**). Together, these observations highlight that the association of IRF6 with patient survival outcomes is highly dependent on the tumor tissue type, pinpointing the complexity of its role and possibly EMT/MET dynamics across different cancers.

**Figure 7.**
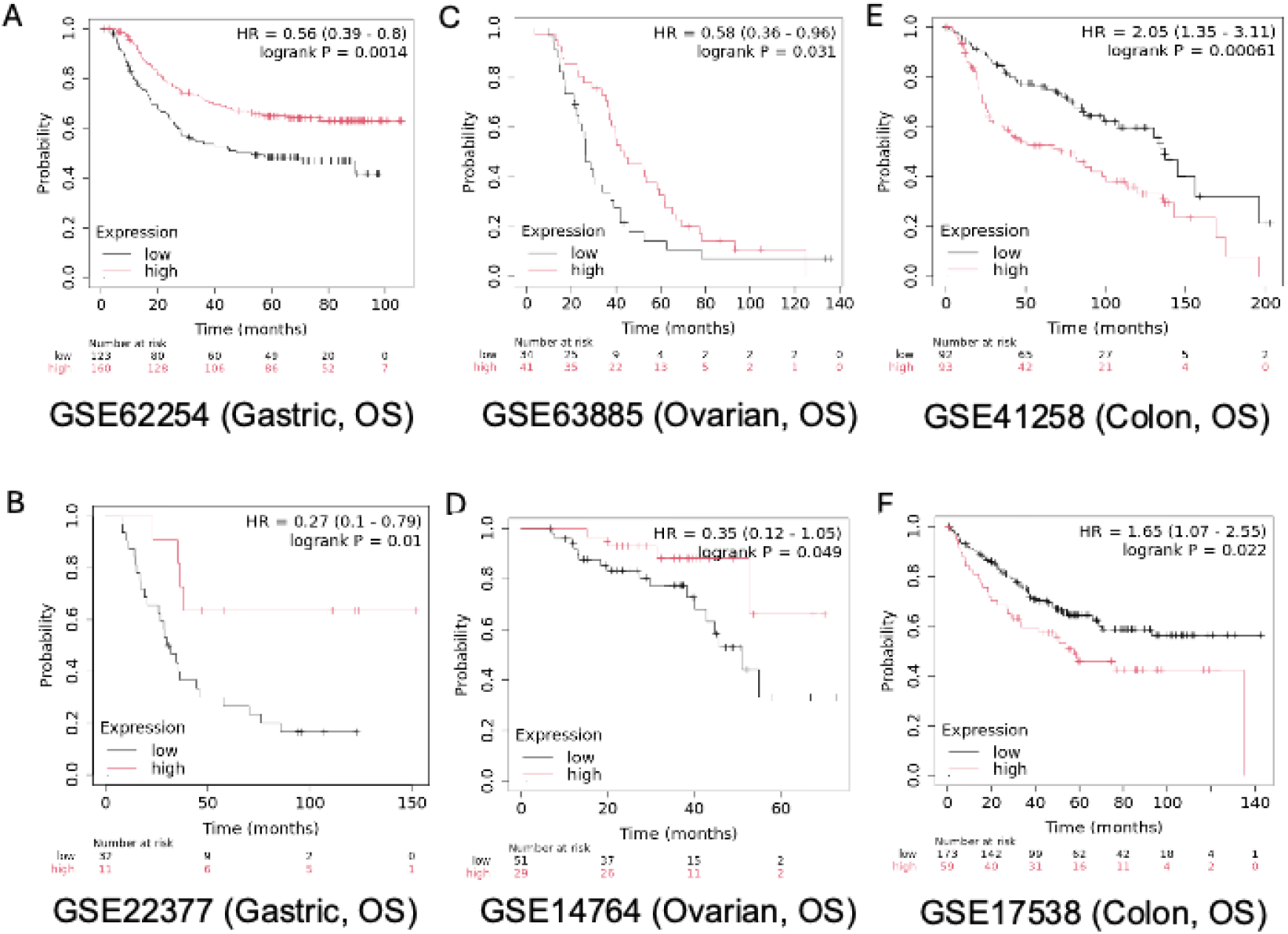
IRF6 correlates with patient survival in a cancer specific manner. A-B) Overall survival trends in GSE62254 and GSE22377 (gastric cancer) respectively. C-D) Overall survival trends in GSE63885 and GSE14764 (ovarian cancer) respectively. E-F) Overall survival trends in GSE41258 and GSE17538 (colon cancer) respectively.

## Discussion

Our integrated findings including transcriptomic data analysis (bulk and spatial), dynamical network modeling (ODE-based and discrete/Boolean modeling) and proof-of-principle experimental validation together suggest IRF6 as an epithelial gatekeeper. These observations are reminiscent of experimental observations that overexpression of IRF6 led to increased levels of many epithelial players (E-cadherin, desmoplakin) and a reduction in mesenchymal markers (Vimentin, N-cadherin) in multiple cancer types. (36,37,72). IRF6 is also known to deliver E-cadherin molecules to plasma membrane of keratinocytes, thus playing key non-transcriptional role in maintaining cell-cell adhesions in the oral epithelium (73,74).

The loss of IRF6, in association with EMT, can also attenuate responses to immunotherapy in pancreatic cancer by promoting resistance to T-cell killing (38). EMT and immunotherapy response are inversely associated through various mechanisms such as, but not limited to, modulation of levels of PD-L1 and other immune checkpoints (75–77). Downregulation of IRF6 by ErbB2 (HER2) can also block anoikis in breast cancer epithelial cells and promote their anchorage-independent growth (78). Anoikis-resistance usually overlaps with a partial EMT (79), suggesting that downregulated IRF6 can poise cancer cells towards a partial EMT, anoikis-resistant and immune-suppressive state. Thus, it is not surprising that IRF6 is downregulated in many carcinomas (36,80) through mechanisms such as DNA methylation (81). Similar downregulation of GRHL3 (80) – a family member of the MET-TF GRHL2 (17,82,83) – is consistent with our observations that IRF6 belongs to the ‘team’ of players inhibiting EMT and/or inducing MET. It is intriguing that IRF6 is likely the only family member among IRFs involved in mediating EMT/MET, unlike many other EMT inhibiting TFs that tend to perform similar or reinforcing functions (OVOL1/2, GRHL2/3, ELF3/5, KLF4/9) (48,84–87).

Besides its role in EMT, IRF6 can also promote sensitivity to cisplatin in colorectal cancer (36), reminiscent of the observation that upregulating KLF4 (belonging to the same ‘team’ as IRF6) can reduce SLUG (88) (a member of the EMT ‘team’) at the protein level and sensitize head and neck cancer cells to cisplatin treatment. Cisplatin resistance has been associated with a partial or complete EMT in multiple cancer types (88,89). Similarly, in pancreatic cancer, IRF6 can enhance susceptibility to T-cell killing by facilitating TNF-induced apoptosis (38). Consistently, in nasopharyngeal carcinoma, IRF6 can repress the ATP-dependent transporter (ABCG2), a protein that confers chemoresistance due to its function as a drug efflux pump (37). Thus, IRF6 re-expression can be a possible strategy to restore sensitivity to many currently used therapies.

We observed that the association of IRF6 with patient survival is tissue-specific. Higher IRF6 expression levels associate with worse survival in colon cancer, but better survival in lung, gastric and ovarian cancer, across many different types of survival (overall, progression-free, relapse-free, post-progression). In renal cell carcinoma and neuroblastoma (90,91), patients with low IRF6 expression had shorter overall and disease-free survival compared to those with high IRF6 expression, but the trend observed was opposite in pancreatic cancer (92). These observations together suggest the need for a comprehensive contextual analysis of the functional roles of IRF6 in different cancers.

Our work contributes to recent efforts to identify regulators of MET such as claudin 7 and C/EBPɑ (93–95). Mapping which nodes are capable of reverting EMT under diverse biochemical (say, genetic background) and biomechanical (e.g., extra-cellular matrix stiffness) conditions is a crucial question to address to suggest strategies that can prevent cells from transitioning to hybrid E/M states (during EMT and/or MET) that can be highly metastatic (96–99). Given the complexity and diversity in cellular transition trajectories during EMT/MET (100,101), systems-level approaches similar to ours would be instrumental in identifying targeting which nodes would lead to successful reprogramming strategies to restrict metastatic burden.

## Author Contributions

MKJ conceptualized research; MKJ and JAS supervised research and obtained funding; ARS, AA, SD, KH and SS performed research and analysed data. All authors contributed to writing and review of the manuscript.

## Data and code availability

All GEO datasets analysed are publicly available, and their accession GSE IDs are mentioned. All the codes used are available here: https://github.com/ARShuba/IRF6

## Materials and Methods

### Mathematical modelling

Coupled ordinary differential equations (ODEs) were used to describe the dynamics of GRN shown in Fig 3A. The level of protein, mRNA or micro-RNA (X) were described by the following typical equation.

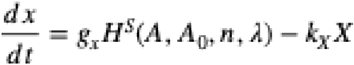

Where g_x_ represents the basal production rate, the shifted Hills function (*H*^*S*^ (*A, A*_0_, *n, λ* ) ) describes the interactions among different molecules, k_x_X describes the first-order degradation of the corresponding molecular species (X). The equations and parameters are available in Supplementary Information (SI). MATLAB (Math-Works Inc.) was used for drawing the bifurcation diagrams using the continuation software package MATCONT (102).

### RACIPE (Random Circuit Perturbation)

Random Circuit Perturbation (RACIPE) was used to investigate the dynamics of GRN shown in Fig 3A over an ensemble of parameters and initial conditions (62). Parameters for solving the ODEs are sampled randomly from within a predefined, biologically relevant range. 10000 parameter sets are sampled and for each parameter set, it initializes each node in the system with a random set of initial conditions (*n* = 100). Using Euler’s method, ODEs are solved to obtain the steady state solutions whose values are then z-normalized and used to perform hierarchical clustering and principal component analysis (PCA).

### Boolean modeling

Boolean modeling is a network topology-centric approach of simulating GRNs. Each node can have two expression levels : -1 and 1, and each edge can have two values : -1 (inhibition) and 1 (activation). The state of the network *S*(*t*) ∈ {−1,1}^*N*^ is a binary vector of length N – the number of nodes in the network, and is updated according to the following update rule (Ising formalism):

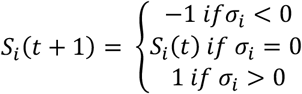

Where *S*_*i*_ (*t*) is the expression of node i at time t, 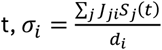 is the threshold function, *J*_*ji*_ is the weight of the edge from node j to node i, and *d*_*i*_ is the indegree of node i.

The network was simulated starting from 100,000 random initial conditions, each initial condition composed of N values sampled uniformly randomly from {-1, 1} Each simulation was performed until a steady state is reached (*S*_*i*_(*t* + 1) = *S*_*i*_ (*t*) ∀ *i*) or 1000 time steps have passed. The updates at each time step was performed asynchronously, by updating one randomly chosen node.

To classify the steady states into their corresponding phenotypes, we devised an E-M score, defined as:

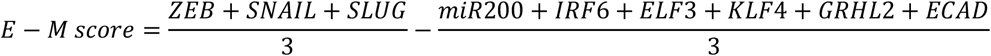

States of E-M score > 1 are classified as mesenchymal, E-M score < -1 as epithelial and remianing states as hybrid phenotypes.

Node knockdown and overexpression experiments were carried out by setting *S*_*i*_ (*t*) = −1 and *S*_*i*_ (*t*) = 1, respectively, for all t. Node deletion experiments were carried out by setting *J*_*ji*_ = 0 ∀ 1 ≤ *j* ≤ *N*.

### Gene expression datasets

GEOquery R Bioconductor package (103) was used to download the gene expression datasets. Gene-wise expression for each sample was obtained by pre-processing the datasets using R.

### siRNA transfection

Cells were seeded onto 12-well glass-bottom plates at 60–70% confluency. Transfection was performed using 10 nM siRNA (final concentration) of IRF6 with RNAiMAX in Opti-MEM and added to cells in complete medium. After 24 hours, the medium was changed, and cells were harvested at 72 hours for immunofluorescence analysis or RNA extraction.

### RNA extraction

For qPCR analysis, total RNA was reverse transcribed using the High-Capacity cDNA Reverse Transcription Kit (Life Technologies). Five-fold diluted cDNA samples were then used for quantitative PCR with KAPA SYBR FAST master mix on the ViiA™7 real-time PCR system (Applied Biosystems). GAPDH mRNA served as the internal control for normalization, and results are expressed as relative expression levels.

### Immunofluorescence microscopy

Cells were seeded at 60-70% confluency in 12-well glass-bottom plates. Briefly, cells were fixed with 4% paraformaldehyde for 15 min at room temperature (RT), permeabilized with PBS containing 0.2% Triton X-100 for 30 min, and blocked with 5% BSA in PBS for 30 min at RT (or overnight at 4 °C). Primary antibodies E-cadherin (1:500) and Zeb-1(1:500) were diluted in 5% BSA in 1XPBS and incubated with cells for 1h at RT or overnight at 4°C. After PBS washes, cells were incubated with anti-rabbit Alexa Fluor 488 secondary antibody (1:2000) and Hoechst (1:2000) for 1h at RT in the dark. Cells were washed, maintained in 1XPBS, and imaged using a fluorescence microscope.

### Single-cell and spatial transcriptomics data analysis

Single-cell RNA-seq data from breast cancer patients (ER+ and triple-negative subtypes – GSE176078) (104) were reanalyzed to evaluate the activity of IRF6-associated (38) and EMT-related transcriptional programs (105). Gene expression matrices were normalized, imputed using MAGIC (106) and pathway activity scores were computed using AUCell (107) based on above mentioned gene signatures. Spatial transcriptomics data from publicly available TNBC patient tissue sections (CID4465, CID44971) (104) were processed using the Seurat pipeline. Expression values were imputed using the MAGIC algorithm prior to signature scoring to enhance gene-gene correlation and pathway resolution.

### Survival analysis

KM-plotter (108) was used for Kaplan–Meier analysis of the different datasets. Sample separation was performed at the median level of gene expression.

