## Supplementary Figures for "Systems-level analysis identifies IRF6 as an inhibitor of epithelial-mesenchymal transition"

### Supplementary Information

#### 1. System of coupled ODEs for mechanistic model

Coupled ordinary differential equations (ODEs) were used to describe the dynamics of the molecular species present in the EMT regulatory circuits shown in Fig 3A. The equations for miR-200, ZEB mRNA, ZEB protein, SNAIL, SLUG mRNA, SLUG, KLF4, ELF3, IRF6, GRHL2 and E-cadherin are correspondingly given as:

$$\frac{d\mu_{200}}{dt} = g_{\mu_{200}} H^s(Z,\lambda_{Z,\mu_{200}}) H^s(S,\lambda_{S,\mu_{200}}) H^s(Sl,\lambda_{Sl,\mu_{200}}) H^s(G,\lambda_{G,\mu_{200}}) - m_Z Y_\mu(\mu_{200}) - m_{Sl} Y_\mu(\mu_{200}) - k_{\mu_{200}}\mu_{200}$$

$$\frac{dm_Z}{dt} = g_{m_Z} H^s(Z,\lambda_{Z,m_Z}) H^s(S,\lambda_{S,m_Z}) H^s(E,\lambda_{E,m_Z}) H^s(G,\lambda_{G,m_Z}) H^s(C,\lambda_{C,m_Z}) - m_Z Y_Z(\mu_{200}) - k_{m_Z} m_Z$$

$$\frac{dZ}{dt} = g_Z m_Z L(\mu_{200}) - k_Z Z$$

$$\frac{dS}{dt} = g_S H^s(I,\lambda_{I,S}) H^s(Sl,\lambda_{Sl,S}) H^s(S,\lambda_{S,S}) H^s(K,\lambda_{K,S}) - k_S S$$

$$\frac{dm_{Sl}}{dt} = g_{m_{Sl}} H^s(S,\lambda_{S,m_{Sl}}) H^s(K,\lambda_{K,m_{Sl}}) H^s(Sl,\lambda_{Sl,m_{Sl}}) - m_{Sl} Y_Z(\mu_{200}) - k_{m_{Sl}} m_{Sl}$$

$$\frac{dSl}{dt} = g_{Sl} m_{Sl} L(\mu_{200}) - k_{Sl} Sl$$

$$\frac{dK}{dt} = g_K H^s(K,\lambda_{K,K}) H^s(Sl,\lambda_{Sl,K}) H^s(S,\lambda_{S,K}) H^s(I,\lambda_{I,K}) - k_K K$$

$$\frac{dE}{dt} = g_E H^s(E,\lambda_{E,E}) H^s(S,\lambda_{S,E}) H^s(Sl,\lambda_{Sl,E}) - k_E E$$

$$\frac{dI}{dt} = g_I H^s(Z,\lambda_{Z,I}) H^s(E,\lambda_{E,I}) H^s(G,\lambda_{G,I}) - k_I I$$

$$\frac{dG}{dt} = g_G H^s(Z,\lambda_{Z,G}) H^s(I,\lambda_{I,G}) - k_G G$$

$$\frac{dC}{dt} = g_C H^s(Z,\lambda_{Z,C}) H^s(G,\lambda_{G,C}) H^s(I,\lambda_{I,C}) - k_C C$$

where, $g_x$ is the innate production rate and $k_x$ is the innate degradation rate, $m_x L(\mu_{200})$ is the net translation rate, $m_x Y_m(\mu_{200})$ is the total mRNA active degradation and $m_x Y\mu(\mu200)$ is the total miR active degradation rate. The shifted hills function $H^S$ is defined as,

$H^S(B,\lambda) = H^-(B)+\lambda H^+(B)$ ,

$H^-(B) = 1/ [1+(B / B_0)^{nB}]$ ,

$H^+(B) = 1-H^-(B)$,

λ is the fold change from the basal synthesis rate due to protein B. λ >1 for activators, while λ<1 for inhibitors.

**2. Parameter Estimation**

We adopted the model parameters for the molecular species constituting the core network (I, miR-200, Snail, Zeb, Slug, KLF4, ELF3) and IRF6, CDH1 and GRHL2 from previously published literature. The parameters are as follows.

| Parameter | Value | Reference |
|---|---|---|
| ks | 0.125 | (Lu et al., 2013) |
| ku200 | 0.05 | (Lu et al., 2013) |
| kmz | 0.5 | (Lu et al., 2013) |
| kz | 0.1 | (Lu et al., 2013) |
| kmsl | 0.5 | Estimated |
| ksl | 0.1155 | (Lu et al., 2013) |
| kk | 0.1732 | (Lu et al., 2013) |
| ke | 0.125 | Estimated |
| ki | 0.154 | Estimated |
| kg | 0.1 | (Jolly et al., 2016) |
| kc | 0.1 | (Mooney et al., 2016) |
| gs | 18000 | (Lu et al., 2013) |
| gu200 | 2100 | (Lu et al., 2013) |
| gmz | 11 | (Lu et al., 2013) |
| gz | 100 | (Lu et al., 2013) |
| gmsl | 90 | Estimated |
| gsl | 50000 | Estimated |
| gk | 50000 | Estimated |
| ge | 50000 | Estimated |
| gi | 5000 | Estimated |
| gg | 200 | (Jolly et al., 2016) |
| gc | 5000 | (Mooney et al., 2016) |
| I0s | 100000 | (Jolly et al., 2017) |
| Z0u200 | 220000 | (Lu et al., 2013) |
| Z0mz | 27500 | (Lu et al., 2013) |
| s0u200 | 180000 | (Lu et al., 2013) |
| s0mz | 180000 | (Lu et al., 2013) |
| u2000 | 10000 | (Lu et al., 2013) |
| sl0u200 | 220000 | Estimated |
| sl0msl | 150000 | Estimated |
| sl0s | 225000 | Estimated |
| s0msl | 180000 | Estimated |
| s0s | 300000 | Estimated |
| k0s | 275000 | Estimated |
| k0msl | 300000 | Estimated |
| s0k | 180000 | Estimated |
| sl0k | 250000 | Estimated |
| k0k | 275000 | Estimated |
| e0e | 200000 | Estimated |

| | | |
|---|---|---|
| sl0e | 220000 | Estimated |
| s0e | 250000 | Estimated |
| e0mz | 180000 | Estimated |
| z0c | 50000 | (Mooney et al., 2016) |
| g0c | 100000 | (Mooney et al., 2016) |
| z0g | 10000 | Estimated |
| i0g | 200000 | Estimated |
| z0i | 80000 | Estimated |
| e0i | 180000 | Estimated |
| g0mz | 25000 | Estimated |
| c0mz | 80000 | Estimated |
| i0k | 180000 | Estimated |
| g0i | 100000 | Estimated |
| g0u200 | 120000 | Estimated |
| i0c | 180000 | Estimated |
| nsmz | 2 | (Lu et al., 2013) |
| nls | 2 | (Jolly et al., 2017) |
| nzu200 | 3 | (Lu et al., 2013) |
| nsu200 | 2 | (Lu et al., 2013) |
| nzmz | 2 | (Lu et al., 2013) |
| nu200 | 6 | (Lu et al., 2013) |
| nslu200 | 1 | (Y. N. Liu et al., 2013) |
| nsls | 3 | (Chen & Gridley, 2013) |
| nsmsl | 1 | (Chen & Gridley, 2013) |
| nss | 5 | (Chen & Gridley, 2013) |
| nks | 2 | (Yori et al., 2011) |
| nsk | 2 | estimated |
| nslk | 4 | (Y.-N. Liu et al., 2012) |
| nkmsl | 2 | (Y.-N. Liu et al., 2012) |
| nkk | 3 | (Mahatan et al., 1999) |
| nee | 3 | (Kopp et al., 2007) |
| nsle | 4 | Estimated |
| nse | 4 | Estimated |
| nemz | 2 | (Suzuki et al., 2021) |
| nzc | 2 | (Mooney et al., 2016) |
| ngc | 3 | (Chung et al., 2016) |
| nzg | 3 | (Jolly et al., 2016) |
| nig | 1 | (De La Garza et al., 2013) |
| nzi | 2 | (D. Li et al., 2019) |
| nei | 5 | (D. Li et al., 2019) |
| ngmz | 1 | (Jolly et al., 2016) |
| ncmz | 2 | (Mooney et al., 2016) |
| nik | 2 | (H. Liu et al., 2016) |
| ngi | 2 | Estimated |

| | | |
|---|---|---|
| ngu200 | 2 | (Chung et al., 2016) |
| nic | 2 | Estimated |
| lamdazu200 | 0.1 | (Lu et al., 2013) |
| lamdasu200 | 0.1 | (Lu et al., 2013) |
| lamdazmz | 7.5 | (Lu et al., 2013) |
| lamdasmz | 10 | (Lu et al., 2013) |
| lamdals | 3 | (Jolly et al., 2017) |
| lamdaslu200 | 0.4 | (Y. N. Liu et al., 2013) |
| lamdasls | 0.5 | (Nakamura et al., 2018) |
| lamdasmsl | 0.5 | (Nakamura et al., 2018) |
| lamdass | 0.4 | (Peiró et al., 2006) |
| lamdakk | 2 | (Dang, 2002) |
| lamdask | 0.25 | (Z. Li et al., 2018) |
| lamdaslk | 0.5 | (Y. N. Liu et al., 2013) |
| lamdakmsl | 0.25 | (Y. N. Liu et al., 2013) |
| lamdaks | 0.5 | (Yori et al., 2011) |
| lamdaee | 4 | (Q. Li et al., 2021) |
| lamdasle | 0.3 | (Lyons et al., 2008) |
| lamdase | 0.25 | (Lyons et al., 2008) |
| lamdaemz | 0.675 | (Suzuki et al., 2021) |
| lamdazc | 0.1 | (Mooney et al., 2016) |
| lamdagc | 3 | (Chung et al., 2016) |
| lamdazg | 0.5 | (Jolly et al., 2016) |
| lamdaig | 5 | (De La Garza et al., 2013) |
| lamdazi | 0.36 | (D. Li et al., 2019) |
| lamdaei | 3 | (D. Li et al., 2019) |
| lamdagmz | 0.1 | (Jolly et al., 2016) |
| lamdacmz | 0.8 | (Mooney et al., 2016) |
| lamdaik | 3 | (H. Liu et al., 2016) |
| lamdagi | 2 | Estimated |
| lamdagu200 | 3 | (Chung et al., 2016) |
| lamdaic | 2 | (Antiguas et al., 2022) |

### Supplementary Figures

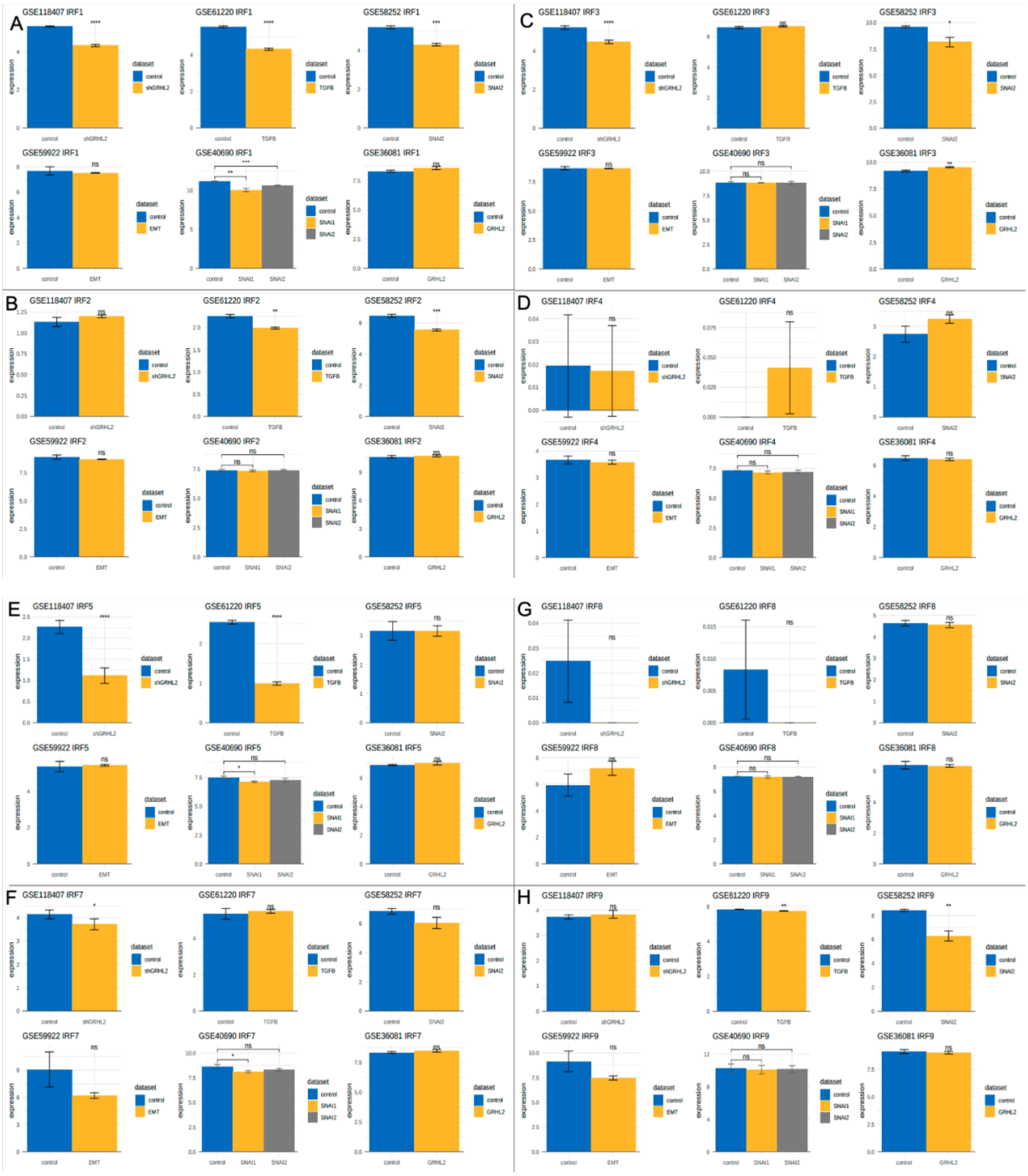

***Fig S1***: *A) Variation in IRF1 expression levels in multiple GEO datasets upon EMT and/or MET induction, corresponding to those shown for IRF6 in Fig 1. B) Same as A) but for IRF2. C) Same as A) but for IRF3. D) Same as A) but for IRF4. E) Same as A) but for IRF5. F) Same as A) but for IRF7. G) Same as A) but for IRF8. H) Same as A) but for IRF9.*

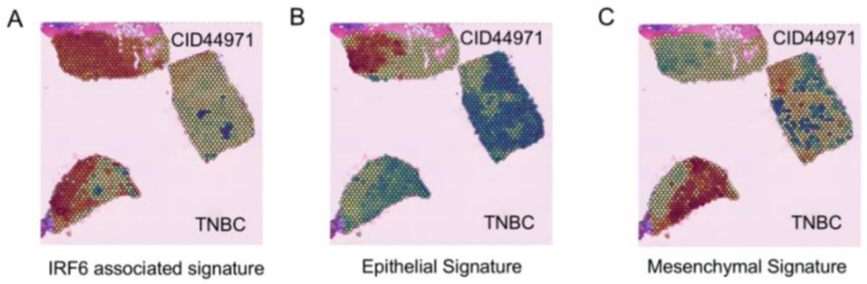


***Fig S2**: Additional spatial transcriptomic views of IRF6, epithelial, and mesenchymal programs in TNBC patient CID44971. Spatial plots of A) IRF6 associated signature, B) breast cancer-specific epithelial signature and C) breast cancer-specific mesenchymal signature activity in TNBC tumor section (CID44971).*

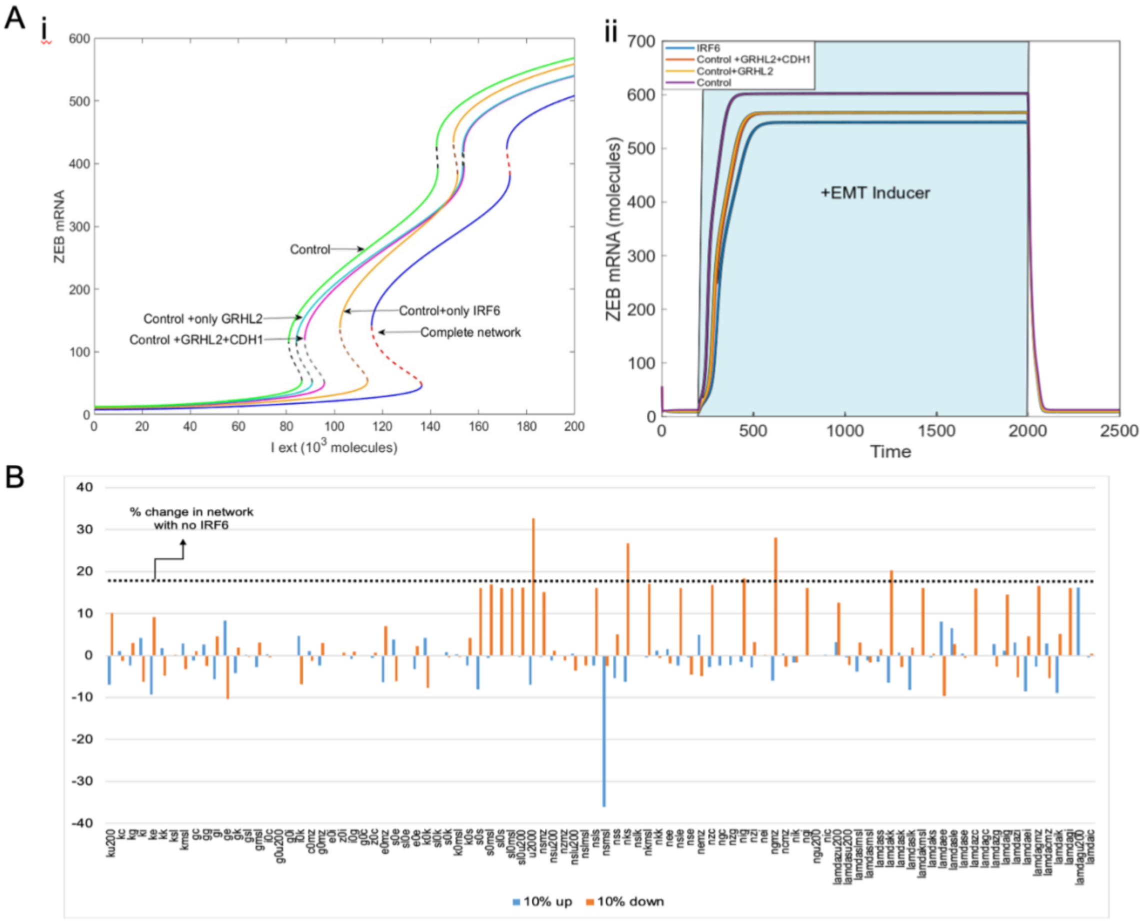


***Fig S3:** A) i) Bifurcation diagram indicating the changing levels of ZEB mRNA levels in response to an external signal (I_ext) level for the core (control) EMT circuit (solid green and dotted black curve), the GRHL2 coupled control circuit (solid blue and dotted black curve), GRHL2 and CDH1 coupled control circuit (solid pink and dotted black curve), only IRF6 coupled control circuit (solid yellow and dotted brown curve) and the IRF6, GRHL2 and CDH1 coupled control circuit (solid blue and dotted red curve). ii) Temporal dynamics of ZEB mRNA levels as a response to a high level of an external EMT signal (I_ext = 200,000 molecules) in a cell of epithelial phenotype (yellow-shaded region) for the circuit shown in Fig 3A. B) Sensitivity analysis indicating percent change in the interval of external signal (I_ext) levels for stable hybrid E/M region, when corresponding parameter values are varied by ±10%. The black dotted line indicates the percent change in the stable hybrid region in the absence of IRF6 (core network+CDH1+GRHL2) when compared to the coupled network with IRF6.*

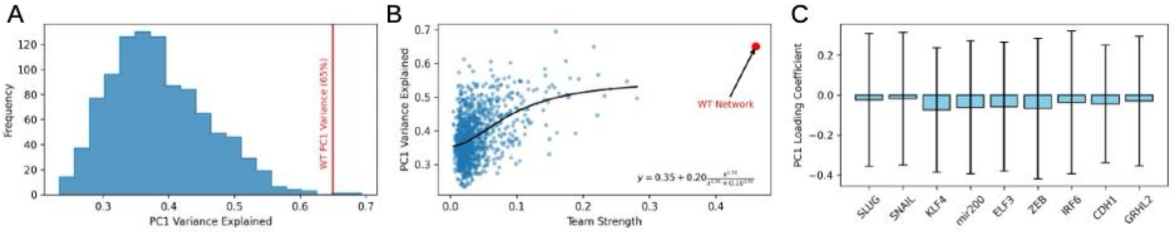


***Fig S4****: A) Histogram showing the distribution of variance explained by PC1 for the 1000 boot-strapped random networks. WT network explains higher variance than most of the random networks. B) Scatter plot showing the PC1 variance vs. team strength (calculated from the influence matrix) for 1000 random networks. C) Plots showing the distribution of values of PC1 loading coefficient observed for each node across the 1000 boot-strapped random networks.*

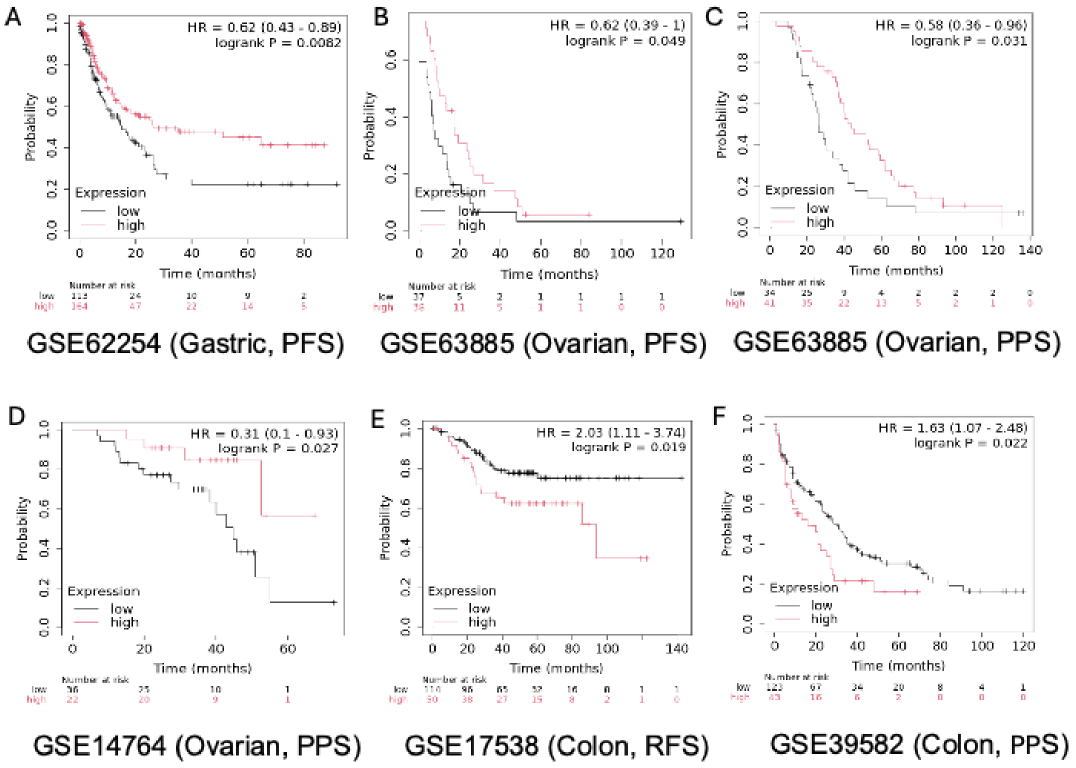


**Fig S5:** *A) Progression free survival trends in GSE62254 (gastric cancer). B-D) Progression-free survival and post-progression survival trends in GSE63885 and GSE14764 (ovarian cancer) respectively. E-F) Relapse free survival and post-progression survival in GSE17538 and GSE 39582 (colon cancer) respectively.*
